# A roadmap for enriching rare cell populations from human post-mortem brain, demonstrated by 100% microglial purity

**DOI:** 10.64898/2026.07.19.739421

**Authors:** Isidora Gocmanac, Hiranyamaya Dash, Maria Weinert, Alexi Nott, Elina Nagaeva, Nathan Skene

## Abstract

Understanding brain disease requires studying the specific cell types that drive each condition: microglia in Alzheimer’s, dopaminergic neurons in Parkinson’s, motor neurons in ALS, and many others. For most of these rare populations, protocols to isolate them from human post-mortem brain at the required purity have never been developed. The shared hurdles are reliable nuclear markers, compatible fluorescent dyes, antibody host-species cross-reactivity, and achieving pure rather than merely enriched populations. Here we present a generalisable roadmap for developing fluorescence-activated nuclear sorting (FANS) protocols for rare cell populations in human brain. We demonstrate it on microglia, reaching 100% purity in cortex and 98% in cerebellum, with the cortex sort simultaneously yielding astrocytes at 93% purity. The roadmap tackles each hurdle including an on-bench antibody labelling step that expands fluorescence channels without cross-reactivity problem and an affordable single-nucleus RNA-seq validation workflow, giving labs a pathway for enriching any rare cell type reliably.

## Introduction

Neurological and psychiatric disorders arise from dysfunction in specific cell types of the brain, and studying them mechanistically therefore requires isolating those cell types from the tissue where the disease actually occurs, most often human post-mortem brain. For that to work, two conditions must be met: the target cell type has to be enriched to high purity, and the enrichment has to be validated. For most rare or disease-relevant populations neither condition is yet routinely satisfied (Fig.1). Sorting protocols exist but have rarely been systematically validated, and purity is often assumed rather than measured.

**Figure 1.**
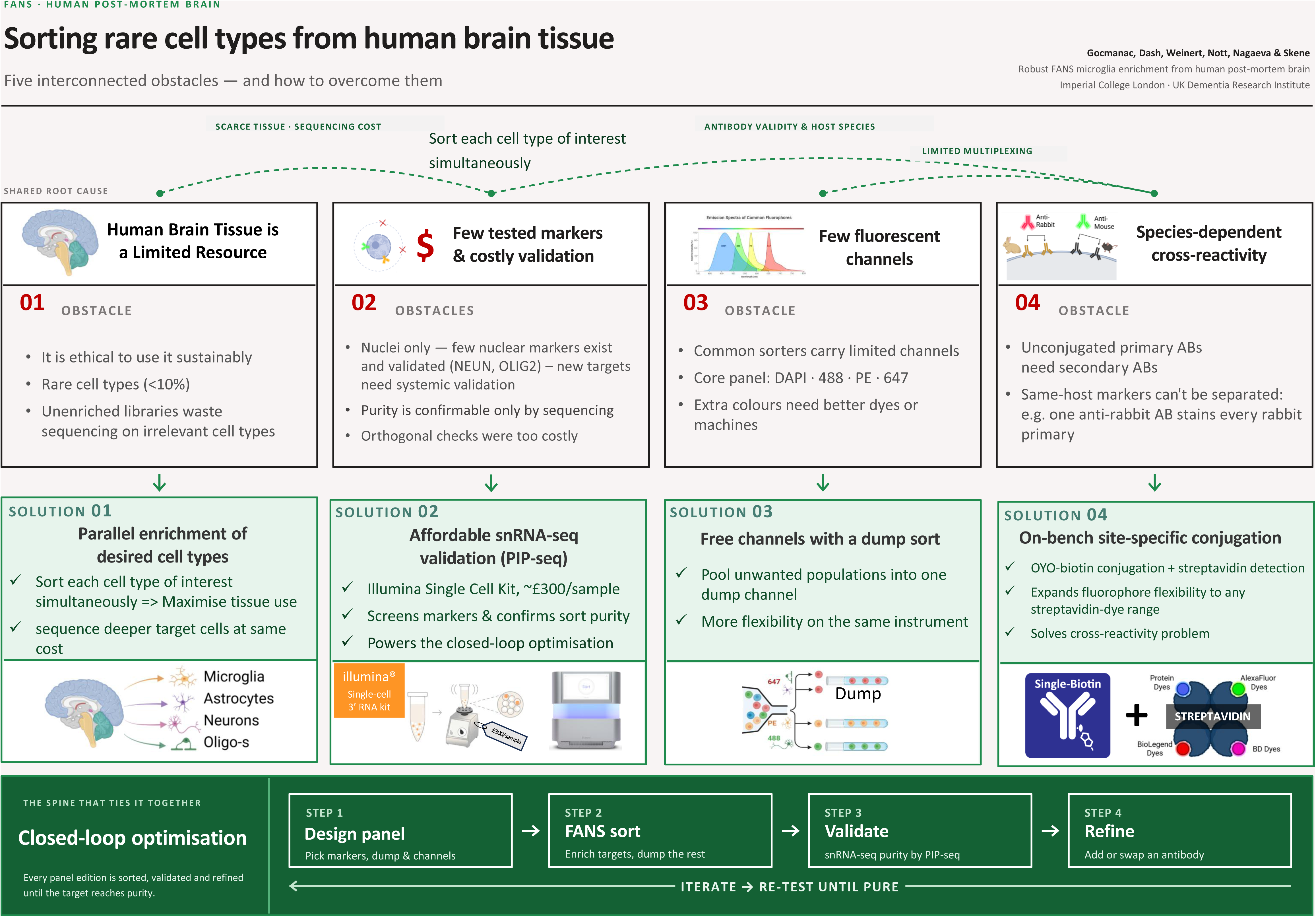
Common obstacles to enriching rare cell types from postmortem human brain tissue, and a roadmap to overcome them. Enrichment of rare cell type populations is constrained by a set of interdependent limitations: tissue availability, marker validation, instrument capacity, and antibody specificity. Rather than addressing these in isolation, our roadmap links each constraint to a practical solution and integrates them into a single closed-loop workflow, in which iterative sorting and sequencing-based validation progressively refine the target population toward purity.

For the major abundant cell classes of the human cortex, sorting works reasonably well: NeuN reliably marks neuronal nuclei^1^ and OLIG2 marks oligodendrocyte-lineage nuclei^2^, and both are routinely used for fluorescent nuclear sorting (FANS) from frozen post-mortem tissue^3^. Beyond these abundant populations, two features of human post-mortem brain constrain what is possible. First, frozen tissue is incompatible with intact-cell recovery: only nuclei can be reliably extracted^4, 5^, which restricts marker choice to nuclear antigens, predominantly transcription factors and structural proteins, rather than the well-validated surface markers routinely used in immunology^6^ (Fig.1). Second, nuclear-antigen antibodies validated for flow cytometry are sparse. Together these constraints make it especially difficult to isolate small populations, such as microglia and specific neuronal subtypes, with the specificity that downstream molecular assays require for reliable read-depth and powered statistical analysis^7, 8^.

Microglia illustrate the problem in its sharpest form. As the resident immune cells of the brain, they are central effectors in Alzheimer’s disease (AD)^9^ and multiple sclerosis (MS)^10^. Genome-wide association studies (GWAS) have mapped the majority of AD common-variant heritability to regulatory elements active specifically in the microglial lineage^11–15^. Yet microglia comprise only 5-10% of human cortical cells^16^, so studies that target them without enrichment systematically under-recover them. Even large single-nucleus RNA-seq efforts in AD deliver only a handful of microglial nuclei per donor and correspondingly few reads per nucleus, leaving these studies under-powered to detect the differentially expressed genes they set out to find^17–20^. Bulk epigenomic assays face the same problem in reverse: unsorted- tissue signal is dominated by more abundant lineages, and any microglia-specific contribution is diluted proportionally^11, 15^. Robust profiling therefore requires upstream enrichment to near- pure (>95%) microglial input (Fig.1).

Microglial nuclear FANS in human post-mortem brain has consequently concentrated on the haematopoietic transcription factor PU.1 (encoded by SPI1)^19, 21^. Recent benchmarking, however, has shown that PU.1⁺ sorts from human cortex reach only ≈60% microglial purity^19^, with the remainder dominated by astrocytes. Notably, these protocols rely on PU.1⁺ positive selection alone, with no prior depletion of the more abundant neuronal, oligodendroglial or astrocytic nuclei, leaving contamination from the more abundant lineages unchecked^19^. IRF8, an alternative microglial transcription factor, has not, to our knowledge, been systematically benchmarked across cortical regions by single-nucleus sequencing^22, 23^. A recent preprint reported >90% purity across several sorted brain cell populations using DNA-methylation deconvolution for validation^24^ but did not explicitly estimate microglial purity by single-cell sequencing. As a result, many laboratories proceed directly from FANS-sorted microglia to bulk epigenomic assays without orthogonal purity validation, because the standard validation method, droplet-based single-nucleus RNA-seq on commercial platforms, costs in the order of £1,500 per sample excluding sequencing, prohibitive at the scale required for routine quality control.

To develop a generalisable roadmap for FANS-panel optimisation of rare cell populations in human post-mortem brain, we used microglia as the target. We describe a five-marker FANS panel, NEUN⁻/OLIG2⁻/IRF8⁺PU.1⁺/SOX9⁻, that pairs three-lineage depletion of neurons (NEUN⁻), oligodendrocytes (OLIG2⁻) and astrocytes (SOX9⁻) with dual IRF8⁺PU.1⁺ positive selection of microglia on a shared channel. The dual-marker strategy ensures microglial capture even when one marker does not label all target nuclei, as we observed for IRF8 in occipital cortex (Fig. 2C). SOX9⁻ exclusion of astrocytes builds on its reported astrocyte- specific nuclear expression in adult brain^25^, which we validate here, to our knowledge for the first time by snRNA-seq on FANS-sorted nuclei from human post-mortem brain. For applications that require additional sorting capabilities despite limited availability of fluorescently conjugated primary antibodies, as in our case for parallel sorting of microglia and astrocytes from a single sample, we further describe an on-bench site-specific OYO-Link biotinylation workflow that circumvents secondary-antibody cross-reactivity. Sort purity is validated by single-nucleus PIP-seq^26^, an orthogonal transcriptomic quality-control workflow approximately 5-fold cheaper than droplet snRNA-seq and cost only £300 per sample excluding sequencing (Fig.1). We benchmark the protocol in occipital cortex and confirm application to parietal cortex and cerebellum, giving laboratories a validated FANS strategy for isolating high-purity microglial nuclei suitable for downstream molecular assays. Importantly, the low-cost optimisation pipeline itself is generalisable to enrichment protocols for other rare cell types in human post-mortem brain (Fig.1).

**Figure 2.**
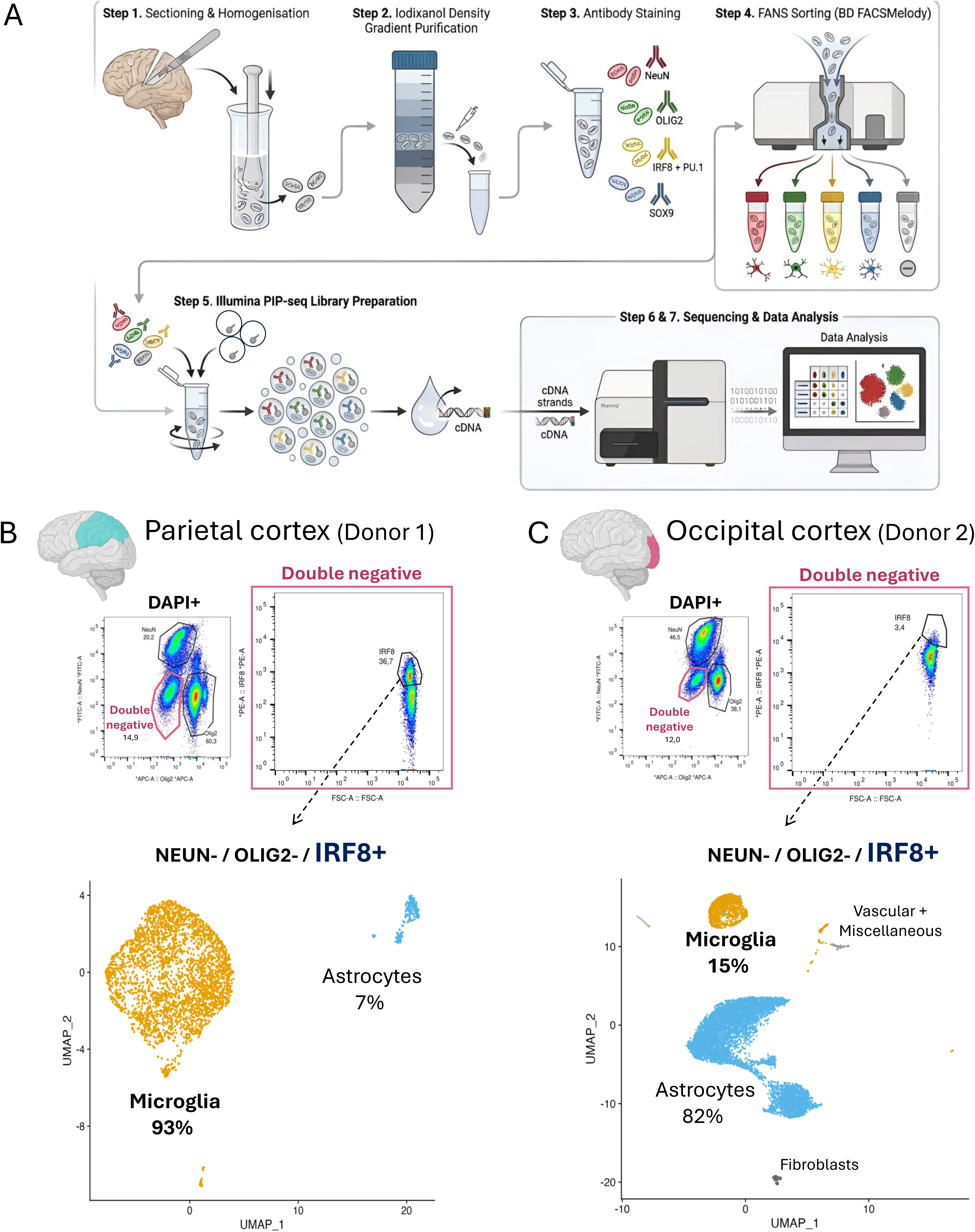
Single-marker IRF8⁺ FANS achieves high microglia purity in parietal but fails in occipital cortex. (A) Schematic of the seven-step FANS–snPIP-seq workflow: cryostat sectioning and dounce homogenisation of frozen post-mortem cortex, iodixanol density-gradient nuclei purification, antibody staining (NEUN, OLIG2, IRF8+PU.1 on a shared channel, SOX9), nuclei sorting on a BD FACSMelody, Illumina PIP-seq library preparation, NovaSeq X Plus sequencing, and computational data analysis. (B) Parietal cortex (Donor 1) IRF8⁺ NEUN⁻ OLIG2⁻ sort (Methods Configuration A) validated by droplet single-nucleus RNA-seq on the 10x Genomics platform: hierarchical FACS gating (DAPI⁺ singlets → NEUN/OLIG2/Double-negative on the AF488 × AF647 plot → IRF8⁺ gate on Double-negative), UMAP embedding of post-sort nuclei coloured by reference- annotated cell type, and donut plot showing cell-type composition (93% microglia, 7% astrocytes; n = 3,106 nuclei). (C) Occipital cortex (Donor 2) IRF8⁺ NEUN⁻ OLIG2⁻ sort with identical layout: FACS gating tree, UMAP and donut showing 15% microglia, 82% astrocytes and ∼3% combined fibroblast / vascular / miscellaneous (n = 8,350 nuclei). Reference cell-type annotations use the Allen Institute MapMyCells workflow against the 10x Whole Human Brain taxonomy (CCN202210140).

## Results

### IRF8⁺ enrichment achieves microglial purity in parietal but fails in occipital cortex

To establish a baseline for nuclear FANS sorting of microglia from human post-mortem cortex, we used IRF8, a microglia-specific transcription factor^22, 23^, to sort IRF8⁺ NEUN⁻ OLIG2⁻ nuclei (Methods Configuration A) from parietal cortex (Donor 1) and occipital cortex (Donor 2), and validated the sorts by droplet-based snRNA-seq on the 10x Genomics platform. In parietal cortex, IRF8⁺ gating yielded a near-pure microglial population: ∼93% microglia and ∼7% astrocytes by reference-anchored MapMyCells annotation (Fig. 2B; Supplementary Fig. 5A). Applied to occipital cortex from a different donor, the same gating strategy recovered only ∼15% microglia and ∼82% astrocytes, with a small (∼3% combined) fibroblast and vascular fraction (Fig. 2C; Supplementary Fig. 5B). Because our two samples came from different regions and different donors, we cannot tell whether this variation is regional, donor-driven, or both, but either way, a more robust protocol is needed. At this point we switched purity validation from 10x Genomics to Illumina single-nucleus PIP- seq (Illumina Single Cell 3’ RNA Prep, T2; Methods), a microfluidics-free, ∼5-fold cheaper alternative that makes routine sort QC tractable, and used it for all subsequent sorts in this study. To reproduce previous poor microglia enrichment with new validation platform, we processed a separate aliquot of the same Donor 2 occipital cortex through the IRF8⁺ NEUN⁻ OLIG2⁻ sort followed by snPIP-seq validation and again recovered an astrocyte-dominated population (∼88% astrocytes, ∼5% neurons; Supplementary Fig. 1). This validated that the IRF8⁺ single-marker protocol does not always enrich pure microglia^24^. As we used two different sequencing platforms (10x Genomics and Illumina snPIP-seq), we believe this is not a sequencing or platform artefact. Inspection of the upstream FANS plots showed that IRF8⁺ versus IRF8⁻ separation within the NEUN⁻ OLIG2⁻ gate was modest in both regions, with overlap between the two populations contributing to gate-placement uncertainty (Fig. 2B, C; Supplementary Fig. 1A). Together, these observations show that single-marker IRF8 sorting cannot perform consistently across cortical regions and donors, and motivated us to optimise the panel in occipital cortex (Donor 2), the more demanding of the two samples tested, for the remainder of the study.

### PU.1⁺ enrichment improves microglial yield in occipital cortex but does not eliminate astrocyte contamination

We next tested PU.1⁺ NEUN⁻ OLIG2⁻ (Methods Configuration A) as an alternative and previously reported^19^ single-marker enrichment strategy in Donor 2 occipital cortex (Fig. 3A). PU.1⁺ sorting recovered substantially more microglia than IRF8⁺ from the equivalent region, with ∼95% of sorted nuclei classifying as microglia and the remaining ∼5% as astrocytes (Fig. 3A; Supplementary Fig. 5C). PU.1 also gave visibly clearer FANS-plot separation between PU.1⁺ and PU.1⁻ populations than IRF8 had in either cortical region, simplifying gate placement and reducing operator-dependent variability. Interestingly, our purity yield overperformed the reported PU.1enrichment efficiency^19^ most likely due to negative sorting of the two most abundant cell types (neurons and oligodendrocytes), but residual astrocyte contamination at the ∼5% level remains incompatible with bulk epigenomic assays that require near-pure input. We therefore reasoned that explicit depletion of astrocytes by negative selection, rather than reliance on positive selection of microglia alone, would be necessary to reach the purity threshold required for cell-type-resolved chromatin profiling.

**Figure 3.**
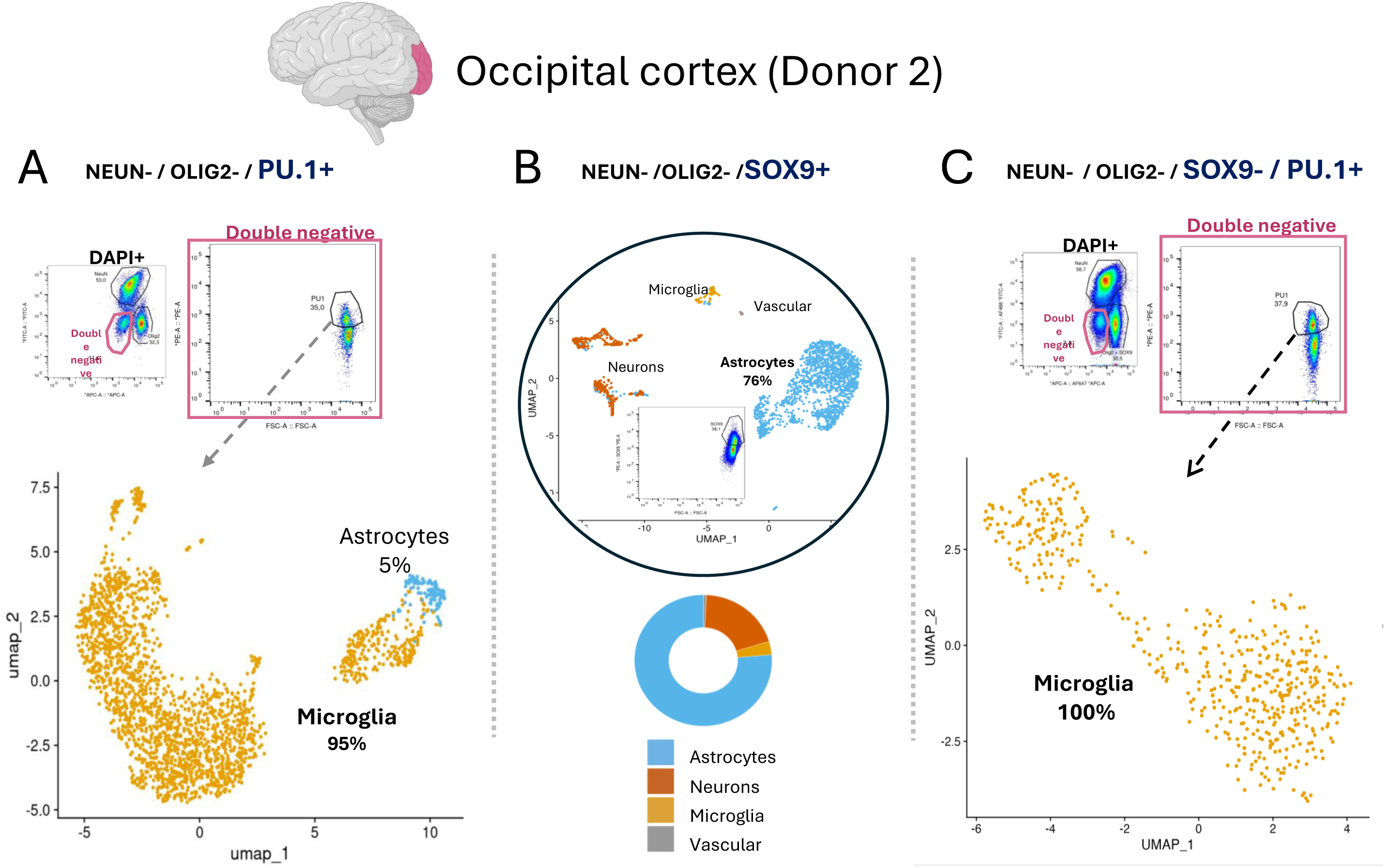
PU.1⁺ enrichment supplemented with SOX9⁻ dump-channel exclusion yields pure microglia in occipital cortex. All sorts performed on Donor 2 occipital cortex and validated by Illumina snPIP-seq. (A) Single-marker PU.1⁺ NEUN⁻ OLIG2⁻ sort (Methods Configuration A): FACS gating (DAPI⁺ → NEUN/OLIG2/Double-negative → PU.1⁺ gate), UMAP embedding and donut plot (95% microglia, 5% astrocytes; n = 2,160 nuclei). (B) Direct SOX9⁺ NEUN⁻ OLIG2⁻ astrocyte enrichment (Methods Configuration B; directly-conjugated anti-SOX9-PE), confirming SOX9 enriches astrocytes (76% astrocytes, 18% interneurons, 3% residual microglia, ∼3% vascular; n = 2,030 nuclei). (C) Combined PU.1⁺ SOX9⁻ NEUN⁻ OLIG2⁻ dump-channel sort demonstrating microglia purity improvement through SOX9-assisted astrocytes removal (Methods Configuration C, PU.1-only variant; OLIG2 and SOX9 co-stained on AF647): FACS gating tree, UMAP and donut showing 100% microglia (n = 520 nuclei).

### SOX9⁺ FANS selectively captures cortical astrocyte nuclei

To enable astrocyte depletion, we first confirmed the specificity of the astrocyte nuclear marker SOX9^25^. Using a directly conjugated anti-SOX9-PE antibody (Methods Configuration B), SOX9⁺ NEUN⁻ OLIG2⁻ nuclei from Donor 2 occipital cortex (Fig. 3B) were sorted and profiled by snPIP-seq. The sorted population was dominated by a large astrocytic cluster (Fig. 3B) in which canonical astrocyte transcripts (GFAP, AQP4, ALDH1L1) were uniformly expressed and microglial and oligodendrocyte markers were largely absent (Supplementary Fig. 2A). Reference-anchored cell-type annotation classified ∼76% of recovered nuclei as astrocytes, with a smaller neuronal fraction (∼18%), residual microglia (∼3%) and rare vascular cells (Fig. 3B). The interneuron contamination is consistent with reported transient SOX9 expression in subsets of cortical interneurons^27^. SOX9 nonetheless remains the most specific nuclear astrocyte marker available, and these data are consistent with earlier flow-cytometry-based validation of SOX9 as a pan-astrocyte nuclear marker in adult brain^25^. To our knowledge this study provides the first orthogonal validation of SOX9-based nuclear FANS in human post-mortem cortex by single-nucleus sequencing. SOX9⁺ selection therefore captures the contaminant we sought to deplete.

### Adding SOX9⁻ astrocyte exclusion to PU.1⁺ selection yields pure microglia population

To reach a pure microglial population, we added SOX9-based astrocyte depletion to the PU.1⁺ enrichment from the previous section. Adding SOX9 to the panel raised two practical problems.

First, secondary-antibody cross-reactivity (Fig.1). Fluorescent secondary antibodies detect their target by binding its host-species constant region, and therefore cannot distinguish between two primary antibodies raised in the same species^6^. Both anti-SOX9 and anti-OLIG2 are raised in rabbit, so an anti-rabbit AF647 secondary used to detect SOX9 also binds the anti-OLIG2-647 even it is a direct conjugate^6^. The two signals therefore necessarily fall into the same fluorescence channel and cannot be resolved separately. Second, fluorescence-channel availability. The panel already occupies all four channels of the standard sorter configuration (DAPI/violet, NEUN-AF488/blue, PU.1-PE/blue, OLIG2-AF647/red). Adding a fifth fluorophore for a distinct SOX9 signal is non-trivial: directly-conjugated anti-SOX9 antibodies in additional channels are rare in compatible host-species/clone combinations, and standard on-bench conjugation kits require high antibody inputs, use non-site-specific chemistry that can compromise antigen binding, and are not cost-justified for the small-scale iterations of panel optimisation. We addressed both problems by co-staining anti-SOX9 and anti-OLIG2 on the same AF647 “dump” channel (Methods Configuration C, PU.1-only variant). Because both antibodies are used here only for negative selection, collapsing their signals into a single exclusion gate loses no sort-relevant information while keeping the panel to three fluorescence channels (NEUN-AF488 depletion, OLIG2+SOX9-AF647 combined depletion, PU.1-PE positive selection). PU.1⁺ SOX9⁻ NEUN⁻ OLIG2⁻ sorting in Donor 2 occipital cortex yielded 100% microglia (n = 520 nuclei; Fig. 3C; Supplementary Fig. 5D), eliminating the residual astrocyte contamination seen with PU.1⁺ enrichment alone (Fig. 3A). Because SOX9 and OLIG2 continue to share the AF647 channel in this configuration, however, the sort still cannot resolve astrocytes as a separate output population, a limitation we address in the next section by moving SOX9 onto its own fluorescence channel.

### The five-marker panel resolves four cortical cell-type populations simultaneously from a single sample

We next wanted to sort all major cortical cell types: neurons, oligodendrocytes, microglia and astrocytes, from a single FANS experiment. The previous panel could not do this because SOX9 and OLIG2 shared the AF647 dump channel, preventing astrocytes from being separated as their own output. To recover astrocytes as a distinct sort, we needed to move SOX9 onto a new fluorescence channel with species-agnostic detection.

To complete the five-marker panel, we (i) added anti-IRF8-PE alongside anti-PU.1-PE to increase the probability of capturing the full microglial population^19^ and improve FANS-plot separation (Fig. 4; Supplementary Fig. 2C); (ii) attached biotin to the anti-SOX9 antibody on- bench using a site-specific OYO-Link photochemical labelling system and detected it with streptavidin-BV786, a dye compatible with the panel and sorter. Because streptavidin binds biotin rather than a species-specific epitope, this detection strategy is immune to the anti- rabbit cross-reactivity that forced SOX9/OLIG2 co-staining in the previous configuration (Methods Configuration D; Supplementary Fig. 2B).

**Figure 4.**
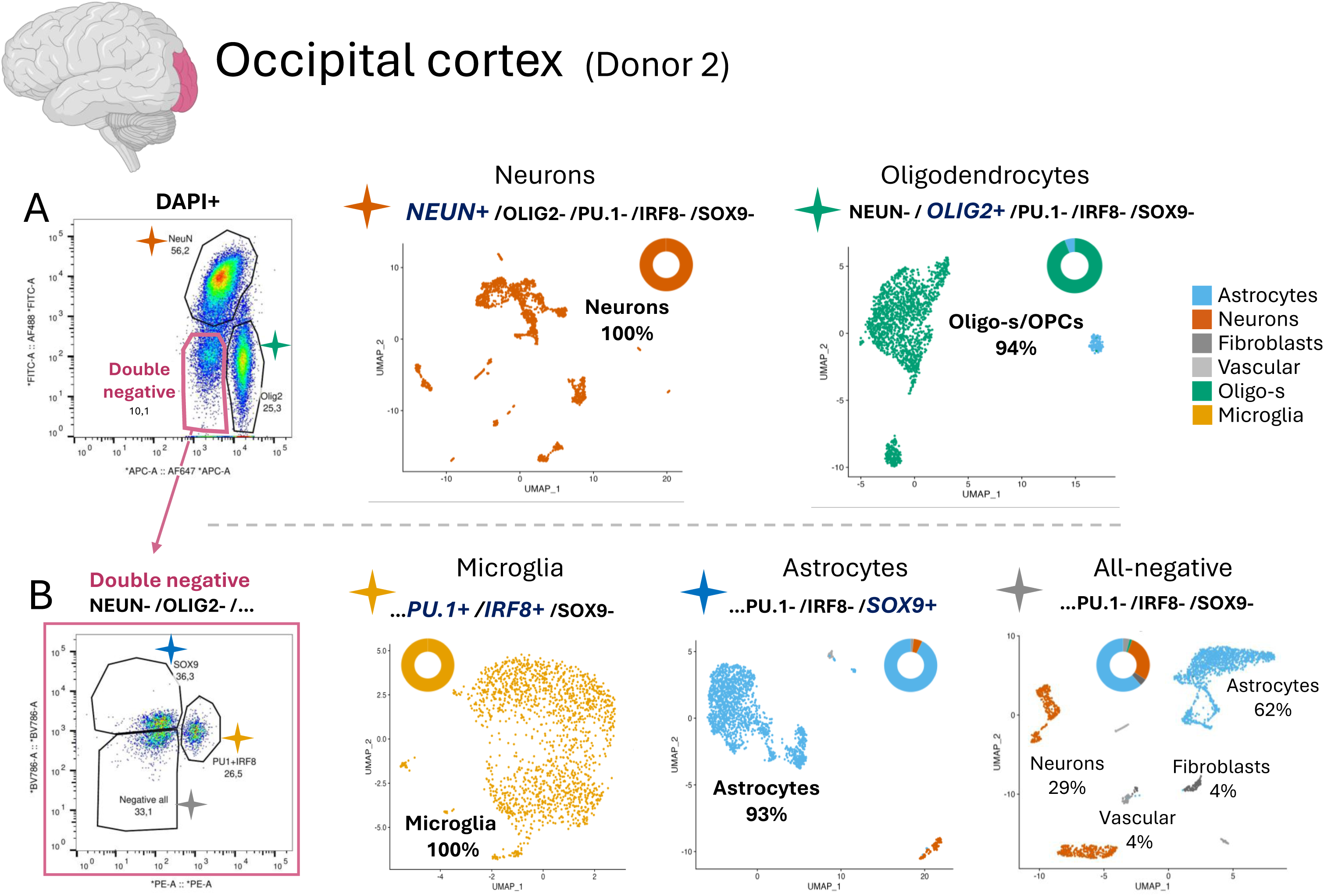
The five-marker panel resolves all major cortical cell types from a single sample. Parallel five-population sort from Donor 2 occipital cortex (Methods Configuration D; OYO-Link biotinylated anti- SOX9 detected with streptavidin-BV786). (A) Lineage gating on the DAPI⁺ AF488 × AF647 plot resolves three populations directly: NEUN⁺ neurons, OLIG2⁺ oligodendrocytes/ OPCs and the NEUN⁻ OLIG2⁻ Double- negative population; UMAP embeddings and donut plots, validated by Illumina snPIP-seq, are shown for the two lineage-positive sorts (NEUN⁺: 100% neurons, n = 1,971; OLIG2⁺: 94% oligodendrocyte lineage, n = 1,841). (B) Sub-gating of the Double-negative population on a BV786 (SOX9) × PE (IRF8+PU.1) plot resolves three further populations: IRF8⁺PU.1⁺ microglia (100% microglia, n = 1,680), SOX9⁺ astrocytes (93% astrocytes, n = 1,747) and an all-negative gate (62% astrocytes, 29% neurons, 4% fibroblast, 4% vascular, 2% oligodendroglial lineage; n = 2,245); UMAP embeddings and donut plots are shown for each, matching star colours link each sorting gate to its corresponding UMAP.

With this configuration we sorted and profiled five gating-defined populations simultaneously from Donor 2 occipital cortex (Fig. 4A, B): NEUN⁺ (neurons), OLIG2⁺ (oligodendrocytes), IRF8⁺ PU.1⁺ (microglia), SOX9⁺ (astrocytes), and the all-negative (NEUN⁻ OLIG2⁻ IRF8⁻ PU.1⁻ SOX9⁻) to characterise its composition and check for any microglial nuclei escaping the positive selection gates. Each sorted population resolved as a distinct UMAP cluster matching its expected cell-type identity by canonical markers and reference annotation (Fig. 4; Supplementary Fig. 3). NEUN⁺ sorting yielded 100% neurons; OLIG2⁺ sorting yielded ∼94% oligodendrocyte lineage (Fig. 4A); SOX9⁺ sorting yielded ∼93% astrocytes; and IRF8⁺ PU.1⁺ sorting yielded 100% microglia (Supplementary Fig. 5E). The all-negative gate was dominated by astrocytes (∼62%) and neurons (∼29%), with smaller fibroblast (∼4%), vascular (∼4%) and oligodendroglial (∼2%) fractions.

To confirm cross-donor and cross-region reproducibility, we repeated the five-population sort on Donor 1 parietal cortex (Fig. 5A) previously used in this project (Fig.2B), recovering 100% neurons (NEUN⁺), 100% oligodendrocyte lineage (OLIG2⁺), 100% microglia (IRF8⁺ PU.1⁺; Supplementary Fig. 5F) and 96% astrocytes (SOX9⁺); the all-negative population contained predominantly 78% astrocytes, 7% neurons, 6% microglia, 2% fibroblast, and 7% oligodendroglial fractions (Supplementary Fig. 4). These results confirm the five-marker panel as a robust cross-donor, cross-region sorting strategy for human cortical nuclei.

**Figure 5.**
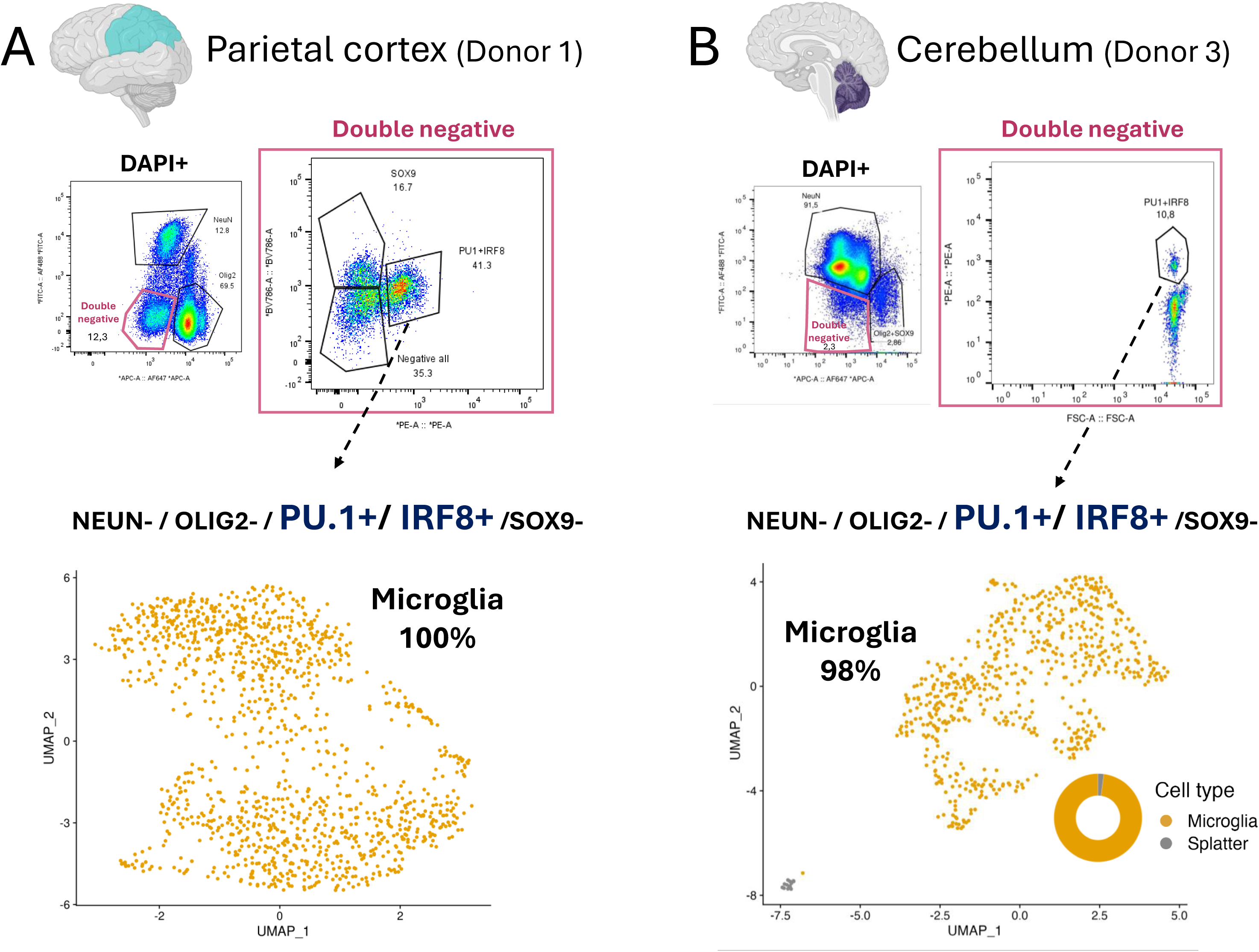
The FANS panel generalises across cortical regions and to cerebellum. (A) Application of the five-marker configuration with OYO-Link biotinylated anti-SOX9 detected via streptavidin- BV786 (Methods Configuration D) to separate Donor 1 parietal cortex aliquot (cf. Fig. 1B). FACS gating tree: DAPI⁺ singlets → NEUN⁻ OLIG2⁻ Double-negative on the AF488 × AF647 plot → SOX9⁻ IRF8⁺PU.1⁺ gate on the BV786 × PE plot. UMAP embedding of post-sort nuclei coloured by reference-annotated cell type, validated by Illumina snPIP-seq: 100% microglia (n = 1,286 nuclei). (B) Application of the dump-channel variant with OLIG2 and SOX9 co-stained on AF647 (Methods Configuration C, IRF8+PU.1 variant) to Donor 3 cerebellum. FACS gating tree: DAPI⁺ singlets → NEUN⁻ OLIG2⁻ SOX9⁻ Double-negative on the AF488 × AF647 plot → IRF8⁺ PU.1⁺ gate on the PE channel. UMAP embedding of post-sort nuclei coloured by reference-annotated cell type and donut plot, validated by Illumina snPIP-seq: 98% microglia (n = 660 nuclei).

### The protocol extends from cortex to cerebellum

Cerebellar tissue differs substantially from cortex in cytoarchitecture, glial composition and neuronal subtype distribution^28^; we therefore asked whether the panel optimised in cortex would generalise to a non-cortical region. Because we did not require parallel sorting of astrocytes from cerebellum, we used the simpler dump-channel variant of the panel (Methods Configuration C, IRF8+PU.1 variant): NEUN-AF488, OLIG2-AF647, IRF8-PE plus PU.1-PE on the microglial-positive channel. Applied to Donor 3 cerebellar tissue, we recovered a microglial population with ∼98% purity (Fig. 5B; Supplementary Fig. 5G), comparable to the values obtained in occipital cortex with the same dump-channel configuration (Fig. 3C). The cells from the residual non-microglial fraction (∼2%) were assigned to splatter neuronal super cluster by MapMyCells^27^. This residual contamination likely reflects the exceptionally high neuronal density of cerebellum^29, 30^ and would require further panel optimisation to eliminate.

## Discussion

Isolating rare cell populations from human post-mortem brain has been a long-standing methodological bottleneck for cell-type-resolved molecular studies of neurological and psychiatric diseases. The roadmap presented here, iterative FANS-panel design coupled to a low-cost snPIP-seq purity read-out, provides a systematic, generalisable route to developing and validating such protocols for any rare nuclear population of interest, on standard flow-cytometry hardware and without dedicated custom conjugation infrastructure (Fig. 1). This work therefore delivers two complementary contributions: a ready-to-deploy protocol for 100% microglial isolation from human cortex, and the generalisable optimisation framework by which any laboratory can build and validate an equivalent sort for other rare cell types.

We demonstrate the efficiency of our roadmap on microglia, the cell type most clearly implicated in AD genetics. The integrated five-marker FANS panel and Illumina snPIP-seq validation workflow reach 100% microglial purity in two cortical donors (Donor 1 parietal and Donor 2 occipital), with portability to cerebellum demonstrated at ≈98% in Donor 3. These purities substantially exceed the ≈60% PU.1⁺ FANS baseline previously reported in human cortex^19^ and ≈90% IRF8⁺ complement DNA-methylation-deconvolution-based estimates^24^ by providing nucleus-resolution single-cell sequencing annotation. The snPIP-seq read-out (≈£300 per sample) makes routine purity quality control tractable at the donor-cohort scales required by genetic-risk studies in AD, MS, and other microglia-involved disorders^9, 10^.

A likely limitation of our approach is that SOX9 does not appear to label the full astrocytic population. Consistent with this, ∼62% of the all-negative gate in the five-marker sort classified as astrocytes (Fig. 4B), suggesting that a subset of astrocytic nuclei escapes SOX9-based capture. This reduces SOX9’s effectiveness as a depletion marker: while incomplete SOX9 labelling did not compromise microglial purity in any of our microglia- targeted configurations, we assume it must have some effect on depletion efficiency overall. For applications aimed at enriching the full astrocytic population rather than depleting it, this limitation matters much more. RFX4 has recently been proposed as an alternative or complementary astrocytic nuclear marker in human brain, with epigenomic data supporting its specificity^31^. We were unable to incorporate RFX4 into our panel systematically, as a stable supply of the verified antibody was not available at the time, whereas SOX9 was readily accessible; combining SOX9 with RFX4 once the RFX4 antibody supply stabilises may help recover the astrocytes that currently fall into the all-negative gate, a possibility worth testing in future work.

Flexible in-lab antibody labelling is a practical necessity for FANS-panel optimisation: sort development requires rapid iteration through candidate antibody-fluorophore combinations, and for most rare nuclear antigens only a small number of clones and directly-conjugated formats are commercially available. On-bench conjugation therefore fills a critical gap, but the chemistries in widest use are not well suited to this application. Most rely on non-site- specific covalent labelling, typically NHS-ester conjugation of surface lysines, which attaches indiscriminately across the antibody, including residues within or adjacent to the Fab paratope, and can therefore reduce antigen-binding activity^32, 33^. The OYO-Link system used here directs photo-chemical conjugation to the Fc region, leaving the antigen-binding site unobstructed^34, 35^; we consider Fc-targeted site-specificity a critical feature for any conjugation chemistry intended for downstream flow-cytometric use.

Three limitations remain. The donor cohort is limited in number (three individuals) and would benefit from expansion, which will inform our understanding of cross-donor reproducibility and disease-state effects. Our snPIP-seq read-out is restricted to cell-type purity validation and is not appropriate for inferring microglial activation states, which require orthogonal confirmation^20^. Finally, while the panel generalised to cerebellum, performance in any new brain region should be validated by snPIP-seq before downstream use and may benefit from region-specific optimisation; for most applications the ≈98% cerebellar purity is already substantially above the threshold required for bulk epigenomic assays.

Together, these results place robust microglia-resolved epigenomic analysis within reach of routine, scalable studies of human post-mortem brain, enabling larger donor cohorts and providing a strong foundation for linking disease-associated regulatory variation to microglial chromatin state without confounding from neighbouring brain cell types. Beyond microglia, we anticipate the roadmap being applied to the many other rare and disease-relevant nuclear populations of human brain that lack validated FANS protocols today.

## Methods

The complete workflow is summarised schematically in Fig. 1A.

### Human post-mortem brain tissue

Frozen post-mortem brain tissue from three neurotypical donors (two male, one female; age range 61–91 years; post-mortem interval 15–48 h; RIN 6.1–7.0) was obtained from the Parkinson’s UK Brain Bank (Imperial College London) under ethics approval Ref. No. 18/WA/0283. Donor 1 contributed parietal cortex, Donor 2 contributed occipital cortex (the region in which the panel was developed and benchmarked), and Donor 3 contributed cerebellum; per-donor demographics, brain bank identifiers, tissue mass and RIN are reported in Supplementary Table 1. All donors were judged free of clinical neurodegenerative disease at death and reported as AD/PD control tissue by the brain bank. Tissue was stored at -80 °C and dissected on dry ice immediately prior to nuclei isolation.

### Nuclei isolation

Iodixanol density-gradient nuclei isolation followed a published protocol^3^ adapted for unfixed human post-mortem brain. Frozen tissue was sectioned on a cryostat (80 µm sections; approximately 300 mg per dissection) onto pre-cooled, RNase-decontaminated surfaces and immediately transferred into pre-cooled Eppendorfs and placed on dry ice. All Eppendorf and FACS tubes were pre-coated overnight with 0.5% BSA in PBS to minimise nuclei loss. Stock solutions, prepared in advance and stored at 4 °C, were 1.5 M sucrose (DNase/RNase-free; Sigma 0335-500G), Iodixanol Medium (IDM: 250 mM sucrose, 150 mM KCl, 30 mM MgCl₂, 60 mM Tris-HCl pH 8.0), 29% and 50% iodixanol working solutions (prepared from IDM and Optiprep (Sigma D1556)) and Nuclei Isolation Medium 1 (NIM1: 250 mM sucrose, 25 mM KCl, 5 mM MgCl₂, 10 mM Tris-HCl pH 8.0). Fresh, on the day of dissection, NIM2 (NIM1 + 1 mM DTT + 1× protease inhibitor cocktail (Promega G6521)) and Homogenisation Buffer (NIM2 + 1 U/µL RNasin RNase inhibitor (Promega N211B) + 0.1% Triton X-100 (Life Technologies HFH10) + 0.1 µg/mL DAPI (Sigma MBD0015)) were prepared on ice. Tissue was dounced in total of 1,000 µL Homogenisation Buffer (HB) in a 2-mL Wheaton glass dounce homogeniser (first, in 800 uL 5 strokes with lose pestle A, then the pestle A was washed with 100 uL HB to avoid nuclei loss, then 10 strokes with tight pestle B, which after was washed with the last 100 uL of HB), with the homogeniser kept on ice throughout. The homogenate was centrifuged at 1,000 × g for 8 min at 4 °C, supernatant discarded, the pellet resuspended in 750 µL HB, and filtered through a 70-µm cell strainer (Falcon 352350). To enrich intact nuclei and deplete myelin, lipid fragments and other low-density debris, the filtrate was mixed 1:1 with 50% iodixanol (final 1,500 µL) and 500 µL aliquots layered onto 500 µL 29% iodixanol cushions in three Eppendorf tubes per sample. Tubes were centrifuged at 5,000 × g for 30 min at 4 °C in a swinging-bucket rotor (as recommended for higher nuclei recovery by 10x best practices https://kb.10xgenomics.com/s/article/360050780051-What-are-the-best-practices-for-working-with-nuclei-samples-for-3-single-cell-gene-expression).

Supernatant was removed from the top, the nuclei pellet resuspended in BSA Buffer (PBS pH 7.4 + 1% MACS BSA (Miltenyi 130-091-376) + 1 U/µL RNasin), pelleted at 500 × g for 5 min at 4 °C, and resuspended in 500 µL BSA Buffer. A 10 µL aliquot was reserved as DAPI- only single-stain control and topped up to 300 µL with BSA buffer; the remaining suspension was used for antibody staining and subsequent sorting.

### Antibody panel and FANS staining

Four configurations of the antibody panel were used across the experiments described above (summarised in Supplementary Table 2), differing in their microglial-marker and SOX9 detection strategies. All four share directly-conjugated anti-NEUN-AF488 (Merck MAB377X, clone A60, mouse; 1:5,000) and anti-OLIG2-AF647 (abcam AB225100, clone EPR2673, rabbit; 1:1,000) as common reagents. Primary/ directly-conjugated antibody staining was performed at 4 °C for 1.5 h in the dark in 500 µL BSA Buffer, followed by a wash in additional 500 µL BSA Buffer (1,000 µL total), pelleted at 500 × g, 5 min, 4 °C and resuspended in 1 mL BSA Buffer for subsequent secondary-antibody or streptavidin staining (where required) at 4 °C for 30 min in the dark, followed by a final wash (centrifugation at 500 × g, 5 min, 4 °C). Nuclei were then resuspended in 500 µL BSA Buffer and filtered through the strainer cap of a 5-mL FACS tube (Falcon 352235) immediately prior to sorting.

**Configuration A:** single-microglial-marker enrichment (Figs. 1B and C, 2A). Anti-NEUN- AF488 (1:5,000), anti-OLIG2-AF647 (1:1,000) and either anti-IRF8-PE (eBioscience 12- 9852-82, clone V3GYWCH, mouse; 1:150; Fig. 1) or anti-PU.1-PE (Cell Signaling 20251S, clone E8I8L, mouse; 1:100). All three antibodies are directly conjugated; no secondary antibody is required. The sort gates IRF8⁺ or PU.1⁺ nuclei.

**Configuration B:** direct SOX9-PE astrocyte validation (Fig. 2B). Anti-NEUN-AF488 (1:5,000), anti-OLIG2-AF647 (1:1,000) and directly-conjugated anti-SOX9-PE (abcam ab224019, rabbit; 1:100). No microglial markers and no secondary antibody are used in this configuration; the sort gates SOX9⁺ nuclei.

**Configuration C:** dual-AF647 dump-channel microglia enrichment (Figs. 2C). Anti-NEUN- AF488 (1:5,000), anti-OLIG2-AF647 (1:1,000), unconjugated rabbit anti-SOX9 (abcam ab225541, azide- and BSA-free formulation; 1:100), and either anti-PU.1-PE alone (1:100; Fig. 2C) or combined with anti-IRF8-PE (1:150; Fig. 4, cerebellum). After primary staining and two washes, nuclei were incubated with donkey anti-rabbit-AF647 secondary antibody (Invitrogen A-31573; 1:1,000). Since the pre-conjugated anti-OLIG2-AF647 is also a rabbit antibody, the anti-rabbit-AF647 secondary will bind it as well; however, this cross-reactivity poses no issue, as both markers serve for negative selection and their combined AF647 fluorescence collectively defines a single dump-channel population.The sort gates IRF8⁺ and/or PU.1⁺ nuclei. SOX9 cannot be sorted as a separate output population in this configuration.

**Configuration D:** five parallel-population panel sort (Fig. 3 and 4A). Anti-NEUN-AF488 (1:5,000), anti-OLIG2-AF647 (1:1,000), anti-IRF8-PE (1:150), anti-PU.1-PE (1:100) and biotinylated rabbit anti-SOX9 (abcam ab225541, azide- and BSA-free formulation; 1:100; biotinylated on-bench using Biotin OYO-Link, Alpha Thera AT4001-100; see “On-bench site- specific OYO-Link biotinylation of anti-SOX9” below) detected with streptavidin-BV786 (BD Biosciences 563858; 1:1,000). Streptavidin-BV786 was applied after the primary-antibody wash, in 1 mL BSA Buffer for 30 min. SOX9 occupies its own fluorescence channel (BV786) in this configuration, enabling parallel sorting of NEUN⁺ neurons, OLIG2⁺ oligodendrocytes, IRF8⁺ PU.1⁺ microglia, SOX9⁺ astrocytes, and an all-negative panel-defined fraction in a single sort.

### On-bench site-specific OYO-Link biotinylation of anti-SOX9

For Fig. 3 and 4A only, rabbit anti-SOX9 (azide- and BSA-free formulation; abcam ab225541) was biotinylated using the Biotin OYO-Link kit (Alpha Thera, AT4001-100) according to the manufacturer’s instructions, immediately before use. The biotinylated antibody was used at 1:100 dilution in BSA Buffer (storage stability beyond same-day use was not assessed, but claimed possible by AlphaThera) and detected with streptavidin-BV786 (BD Biosciences 563858; 1:1,000).

### FANS sorting and gating strategy

Sorting was performed on a BD FACSMelody cell sorter at the Imperial College London Flow Cytometry Facility, equipped with 405 nm, 488 nm and 640 nm lasers (PE excited at 488 nm via cross-excitation) and operated with a 100 µm nozzle. Spectral compensation was applied using the BD FACSMelody’s automated compensation tool with channel-specific defaults for DAPI, AF488, PE, AF647 and BV786, supplemented with single-stain compensation beads where required. A DAPI-only control (no antibodies) was prepared in parallel for every sort to anchor the singlet/DAPI⁺ gate. For Configuration D, a no-primary negative control (streptavidin-BV786 added without biotinylated anti-SOX9) was prepared in parallel to confirm that BV786 signal was specific to biotinylated SOX9 (Supplementary Fig.2B). Stained nuclei were filtered through the strainer cap of a 5-mL FACS tube immediately before sorting and gated hierarchically (Supplementary Fig. 6): FSC-A × SSC-A nuclei → SSC-A × SSC-W singlets → DAPI⁺ on DAPI-A × FSC-A → AF488 × AF647 lineage plot → microglial sub- gating out of Double Negative population on a PE-A × FSC-A plot (Configurations A–C) or on a BV786 × PE plot (Configuration D). On the AF488 × AF647 plot, Configurations A–C draw only the NEUN⁻ OLIG2⁻ double-negative gate; Configuration D additionally gates NEUN⁺ neurons and OLIG2⁺ oligodendrocytes for parallel collection. In Configuration D, the downstream BV786 × PE plot resolves PE⁺ BV786⁻ microglia, BV786⁺ PE⁻ astrocytes, and an all-negative panel-defined fraction. Sorted nuclei were collected into BSA Buffer supplemented to 0.8 U/µL RNasin and 1× protease inhibitor cocktail in pre-coated DNA- LoBind tubes (Eppendorf 22431005) maintained at 4 °C, and counted on a Luna fluorescence cell counter (Logos Biosystems) with AO/PI viability stain (Logos Biosystems F23001).

### Single-nucleus RNA-seq for FANS validation

Sorted nuclei were validated by single-nucleus RNA-seq using two platforms. For the initial benchmarking experiments (Fig. 1B and C), nuclei were processed with the Chromium Next GEM Single Cell 3′ v3.1 chemistry (10x Genomics, PN-1000268) following manufacturer instructions for nuclei input. For all subsequent experiments, nuclei were processed with the Illumina Single Cell 3′ RNA Prep, T2 kit (Illumina catalogue 20135689), which uses Particle- templated Instant Partitions (PIP-seq) chemistry developed by Fluent BioSciences. The kit supports cell or nuclei input; for nuclei the lysis temperature is 66 °C (versus 37 °C for cells). Nuclei were loaded at 1,250 nuclei/µl in 5 µl per reaction (target input 5,000 nuclei) together with 40 U (1 uL) RNase inhibitor in a final loading volume of ≤ 6 µl, per the manufacturer’s nuclei protocol. Capture, lysis, mRNA isolation, reverse transcription, cDNA amplification, library fragmentation, adapter ligation and library amplification were performed according to manufacturer instructions. The transition from 10x to PIP-seq for routine validation was motivated by cost: at our institutional sequencing core, 10x Genomics 3’ v3.1 library preparation costs approximately £1,500 per sample + sequencing cost, compared with approximately £300 per sample for PIP-seq + sequencing cost, making validation of every FANS sort economically tractable. Both platforms generate 3’-tagged single-nucleus libraries that yield comparable cell-type composition estimates after standard read alignment, count- matrix generation and clustering.

### Sequencing

Concentration and size distribution of final libraries were assessed by Qubit dsDNA HS and TapeStation HSD5000. Sequencing was performed on a NovaSeq X Plus using paired-end PE150 reads. Target sequencing depth was 50,000 read pairs per recovered nucleus.

### Computational analysis

Sequencing results were uploaded to Illumina BaseSpace Sequence Hub^36^. Samples with identifiers SK493_* were processed using 10x Genomics Cell Ranger (version 9.0.1) with the *refdata-gex-GRCh38-2024-A* (10x Genomics standard) reference. All other samples were processed using the Illumina DRAGEN single-cell RNA-seq pipeline (version 4.4.6)^37^ with the *hg38_alt_masked_v5* reference genome. The resulting filtered feature-barcode matrices were used for downstream analyses.

Gene-barcode matrices were imported into R (version 4.5)^38^ using Seurat (version 5.5.0)^39–43^. Cells were filtered to retain barcodes with at least 200 detected RNA features and no more than 10% mitochondrial UMIs. Doublets were identified independently for each sample using DoubletFinder (version 2.0.6)^44^ and removed prior to downstream analysis.

Samples were normalised, scaled, clustered and embedded independently using Seurat with 2,000 variable features, 25 principal components, graph-based clustering and UMAP dimensionality reduction. Technical batch pairs in the test cohort were integrated using Seurat CCA integration. No technical batch integration was performed in the final cohort.

Post-QC raw count matrices were exported as H5AD files and annotated using the Allen Institute MapMyCells^45^ workflow with *hierarchical mapping* against the *10x Whole Human Brain reference taxonomy (CCN202210140)*. Returned annotations were imported into the Seurat object as metadata. Cell type labels correspond to the MapMyCells supercluster annotation within each Seurat cluster. Neuronal superclusters were then manually collapsed into “Neurons”, and oligodendrocyte and OPC superclusters were collapsed into “Oligo-s” for the same colour representation. For the Parietal Cortex sample Donor 1, cells assigned by the MapMyCells as Miscellaneous supercluster were collapsed into “Neurons”, as they expressed a range of canonical neuronal markers (Supplementary Fig.4A).

### Statistics and reproducibility

No statistical method was used to predetermine sample size. All cells passing predefined QC and doublet-removal criteria were included in the analysis. Analyses were descriptive unless otherwise stated. Randomised steps were made reproducible using fixed seeds for DoubletFinder and UMAP. The complete workflow was implemented as ordered R scripts with intermediate checkpoints and tabulated QC summaries, enabling reproducible reruns from the DRAGEN output matrices.

## Data availability

Processed DRAGEN output matrices are available through the NCBI Gene Expression Omnibus (GEO) under accession *GSE333448*. The matrices used for downstream analyses correspond to the DRAGEN-generated *filtered_feature_bc_matrix* outputs from 10x Genomics Cell Ranger and Illumina BaseSpace. The raw sequencing data are available upon request.

## Code availability

Analysis code is available at https://github.com/neurogenomics/microglia-sorting. The repository expects DRAGEN/10x-compatible matrices under *data/matrices/<cohort>/<sample>/filtered_feature_bc_matrix/*. Downstream checkpoints, figures and tables can be reproduced by running the numbered R scripts in *scripts/test/* or *scripts/final/*.

Matrices can be obtained from NCBI GEO as described in the Data Availability section. Additional setup instructions are provided in the repository README.

## AI usage disclosure

AI was used to help implement and document the R/Seurat analysis scripts for single-cell QC, clustering, cell-type annotation, and figure generation. The overall analytical design and code architecture were conceived by the authors and all AI-assisted code and outputs were reviewed and validated manually. AI was also used to proofread the manuscript and improve language quality. Summary Figure 1A was generated in FigureLabs platform (www.figurelabs.ai).

## Acknowledgements

The authors would like to acknowledge the Parkinson’s UK Brain Bank, funded by Parkinson’s UK (a registered charity in England and Wales, No. 258197, and in Scotland, SC037554), for supplying the human tissue samples and associated clinical/neuropathological data used in this study. This work has been supported by the Imperial College Flow Cytometry Facility, which is infrastructure funded by the NIHR Imperial Biomedical Research Centre (BRC); we particularly thank Dr Joana Carrelha and Fredrik Wallberg for expert assistance with FANS sorting on the BD FACSMelody. We thank Dr Daria Gavriouchkina and Modesta Blunskyte of the UK DRI Single-Cell and Spatial Omics (SCSO) Facility at University College London for assistance with 10x Genomics library preparation for the two cortical benchmarking samples; NovaSeq X Plus sequencing was performed by Novogene. We are especially grateful to Marianna Papageorgopoulou (Paul Matthews laboratory, UK DRI at Imperial College London) for training I.G. in the iodixanol density-gradient nuclei isolation protocol that forms the methodological foundation of this work. We thank members of the Nott laboratory for generously sharing reagents at key points during the project. This work is supported by the UK Dementia Research Institute award number UK DRI-5008 through UK DRI Ltd, principally funded by the UK Medical Research Council. N.S. also received funding from a UKRI Future Leaders Fellowship (grant number MR/T04327X/1).

## Supplementary tables

**Supplementary Table 1.** Donor metadata. Per-donor demographics, brain bank identifiers, brain region used, tissue mass, RIN and neuropathology score for the three control donors used in this study. Tissue was obtained from the Parkinson’s UK Brain Bank (Imperial College London).

| Donor | Brain bank ID | Sex | Age (years) | PMI (h) | Brain region | RIN | Braak / CERAD |
| --- | --- | --- | --- | --- | --- | --- | --- |
| 1 | PDC029 | M | 82 | 48 | Parietal cortex | 6.5 | Not formally staged; control |
| 2 | PDC040 | F | 61 | 15 | Occipital cortex | 6.1 | Not formally staged; control |
| 3 | PDC078 | M | 91 | 18 | Cerebellum | 7.0 | Braak 2; CERAD neg; Thal Aβ 2 |
| 3 | PDC078 | M | 91 | 18 | Parietal cortex | 5.5 | Braak 2; CERAD neg; Thal Aβ 2 |

**Supplementary Table 2.** Antibodies and reagents. All antibodies used in this study, with the panel configurations (A–D, see Methods) in which each is used. Em- dash indicates not applicable (e.g., for non-antibody reagents or directly-conjugated antibodies for which clone or host is not specified by the supplier).

| Antibody / reagent | Clone | Host | Conjugate | Supplier | Catalog | Dilution | Configurat ions |
| --- | --- | --- | --- | --- | --- | --- | --- |
| anti-NEUN | A60 | Mouse | AF488 | Merck | MAB377X | 1:5,000 | A, B, C, D |
| anti-OLIG2 | EPR2673 | Rabbit | AF647 | abcam | AB225100 | 1:1,000 | A, B, C, D |
| anti-IRF8 | V3GYWCH | Mouse | PE | eBioscience | 12-9852-82 | 1:150 | A, C, D |
| anti-PU.1 | E8I8L | Mouse | PE | Cell Signaling | 20251S | 1:100 | A, C, D |
| anti-SOX9 (preconjugated) | EPR14335-78 | Rabbit | PE | abcam | ab224019 | 1:100 | B |
| anti-SOX9 (BSA/azide-free) | EPR14335-78 | Rabbit | unconjugated | abcam | ab225541 | 1:100 | C, D |
| Donkey anti-rabbit | — | Donkey | AF647 | Invitrogen | A-31573 | 1:1,000 | C |
| Streptavidin-BV786 | — | — | BV786 | BD Biosciences | 563858 | 1:1,000 | D |
| Biotin OYO-Link | — | — | — | Alpha Thera | AT4001-100 | (reagent) | D |
| DAPI | — | — | — | Sigma | MBD0015 | 0.1 µg/mL | All (nuclei stain) |

## Supplementary figures

**Supplementary Figure 1. Cross-platform validation of the IRF8⁺ failure in occipital cortex.** (A) Repeat of the IRF8⁺ NEUN⁻ OLIG2⁻ sort on a separate Donor 2 occipital cortex aliquot (Methods Configuration A) processed by Illumina snPIP-seq library preparation rather than 10x Genomics (cf. Fig. 1C): hierarchical FACS gating tree (DAPI⁺ → NEUN/OLIG2/Double-negative → IRF8⁺ gate). (B) UMAP embedding and donut plot showing 88% astrocytes, 5% microglia and ∼7% other (n = 760 nuclei). The reproducible recovery of an astrocyte-dominated population across two orthogonal sequencing platforms confirms that the IRF8⁺ failure in occipital cortex reflects an intrinsic marker–region and/or donor limitation rather than a platform artefact.

**Supplementary Figure 2. Validation of cell-type specificity, detection chemistry and dual-marker selection underlying the five-marker FANS panel.**

(A) Feature plots showing expression of canonical cell-type markers in the SOX9-PE enriched sample from Donor 2 occipital cortex: CX3CR1, P2RY12, AIF1 (microglia); GFAP, AQP4, SOX9 (astrocytes); RBFOX3, SNAP25 (neurons); OLIG1, OLIG2 (oligodendrocytes); DCN (fibroblast); CLDN5 (vascular). Expression values are log-normalised and all plots share the same scale. (B) Validation of the BV786-streptavidin detection system used for SOX9 in Methods Configuration D (Donor 3 parietal cortex): three side-by-side BV786-A versus PE-A scatter plots showing, from left to right, the negative control (sample stained without anti-SOX9 primary; SOX9- biotin⁻ / Streptavidin-BV786⁺), the stained sample (SOX9-biotin⁺ / Streptavidin-BV786⁺), in which a discrete SOX9⁺ population is present, and BV786 control beads (instrument positive control). The selective appearance of the SOX9⁺ population in the stained sample only confirms the specificity of the biotinylated anti-SOX9 / streptavidin-BV786 detection chain. (C) Comparison of IRF8-PE alone versus combined IRF8-PE + PU.1-PE staining on Donor 2 occipital cortex on the PE channel; the right plot shows improved separation of the PE⁺ (microglia) and PE⁻ (Triple-negative) populations with dual-marker selection on a shared PE channel.

**Supplementary Figure 3. Canonical cell-type marker expression across the five sorted populations from the Configuration D five-marker sort (Donor 2 occipital cortex).**

Feature plots showing expression of canonical cell-type markers: CX3CR1, P2RY12, AIF1 (microglia); GFAP, AQP4, SOX9 (astrocytes); RBFOX3, SNAP25 (neurons); OLIG1, OLIG2 (oligodendrocytes); DCN (fibroblast); CLDN5 (vascular), on the UMAP embedding of each sorted population from Configuration D: (A) NEUN⁺ neurons; (B) OLIG2⁺ oligodendrocytes; (C) IRF8⁺PU.1⁺ microglia; (D) SOX9⁺ astrocytes; (E) all-negative gate. Marker-expression patterns are consistent with the reference-anchored cell-type annotations reported in Figure 3. Expression values are log-normalised and all plots within each sorted population share the same scale.

**Supplementary Figure 4. Cross-region replication of the five-marker panel in Donor 1 parietal cortex.**

Parallel five-population sort, repeated for another donor and cortical region (cf. Fig. 3, Supplementary Fig.3), reproduces the efficiency of the five-marker panel (Methods Configuration D; OYO-Link biotinylated anti-SOX9 detected with streptavidin-BV786). Feature plots showing expression of canonical cel l-type markers: CX3CR1, P2RY12, AIF1 (microglia); GFAP, AQP4, SOX9 (astrocytes); RBFOX3, SNAP25 (neurons); OLIG1, OLIG2 (oligodendrocytes); DCN (fibroblast); CLDN5 (vascular), on the UMAP embedding of each sorted population from Configuration D applied to Donor 1 parietal cortex: (A) NEUN⁺ neurons (100%; n=3,803); (B) OLIG2⁺ oligodendrocytes/OPCs (100%; n=1,963); (C) IRF8⁺PU.1⁺ microglia (100%; n=1,286); (D) SOX9⁺ astrocytes (96% astrocytes, 4% neurons; n= 1,208); (E) all-negative gate (77% astrocytes, 7% oligodendrocytes, 6% microglia, 4% neurons, 3% miscellaneous, 2% fibroblasts; n=1,450). Marker-expression patterns confirm the reference-anchored cell-type annotations depicted at the associated UMAPs. Expression values are log-normalised and all plots within each sorted population share the same scale.

**Supplementary Figure 5. Microglia subtype-marker expression across all microglia-enriched sorts.**

Feature plots showing expression of microglial homeostatic and activation-associated markers on the UMAP embedding of each microglia-targeted sort in the study: (A) Donor 1 parietal cortex IRF8⁺ NEUN⁻ OLIG2⁻ (Methods Configuration A, 10x Genomics); (B) Donor 2 occipital cortex IRF8⁺ NEUN⁻ OLIG2⁻ (Configuration A, 10x Genomics); (C) Donor 2 occipital cortex PU.1⁺ NEUN⁻ OLIG2⁻ (Configuration A, snPIP-seq); (D) Donor 2 occipital cortex PU.1⁺ SOX9⁻ NEUN⁻ OLIG2⁻ (Configuration C, PU.1-only variant, snPIP-seq); (E) Donor 2 occipital cortex IRF8⁺PU.1⁺ five-marker sort (Configuration D, snPIP-seq); (F) Donor 1 parietal cortex IRF8⁺PU.1⁺ five-marker sort (Configuration D, snPIP-seq); (G) Donor 3 cerebellum IRF8⁺PU.1⁺ NEUN⁻ OLIG2⁻ SOX9⁻ (Configuration C, IRF8+PU.1 variant, snPIP-seq). Expression values are log-normalised and all plots within each sorted population share the same scale.

**Supplementary Figure 6. Hierarchical FANS gating strategy.**

Representative gating from a Donor 2 occipital cortex sort following Methods Configuration D. Plots are shown from left to right in the order of the gating hierarchy: (i) FSC-A × SSC-A “Nuclei” gate (83.3% of acquired events) excluding debris and aggregates; (ii) SSC-W × SSC-A “Singlets” gate (97.5%) excluding doublets; (iii) DAPI-A × FSC-A “DAPI⁺” gate (83.4%) selecting intact nuclei from cellular debris; (iv) AF488 (NEUN) × AF647 (OLIG2) lineage plot resolving NEUN⁺ neurons (54.2%), OLIG2⁺ oligodendrocytes (29.3%) and the NEUN⁻ OLIG2⁻ “Double-negative” population (10.6%); (v) BV786 (SOX9) × PE (IRF8+PU.1) sub-gating of the Double-negative population into SOX9⁺ astrocytes (39.1%), IRF8⁺PU.1⁺ microglia (25.2%) and an all-negative panel-defined fraction (33.5%). In Configurations A–C, the first four gate strategies are identical to those shown here, but the BV786 × PE sub-gating step (rightmost plot) is replaced by a PE-A × FSC-A plot used to draw the microglial- positive gate on the Double-negative population (see Methods).

## Notes

### Competing Interest Statement

The authors have declared no competing interest.

