## Supplemental Figures 1-6 for "A roadmap for enriching rare cell populations from human post-mortem brain, demonstrated by 100% microglial purity"

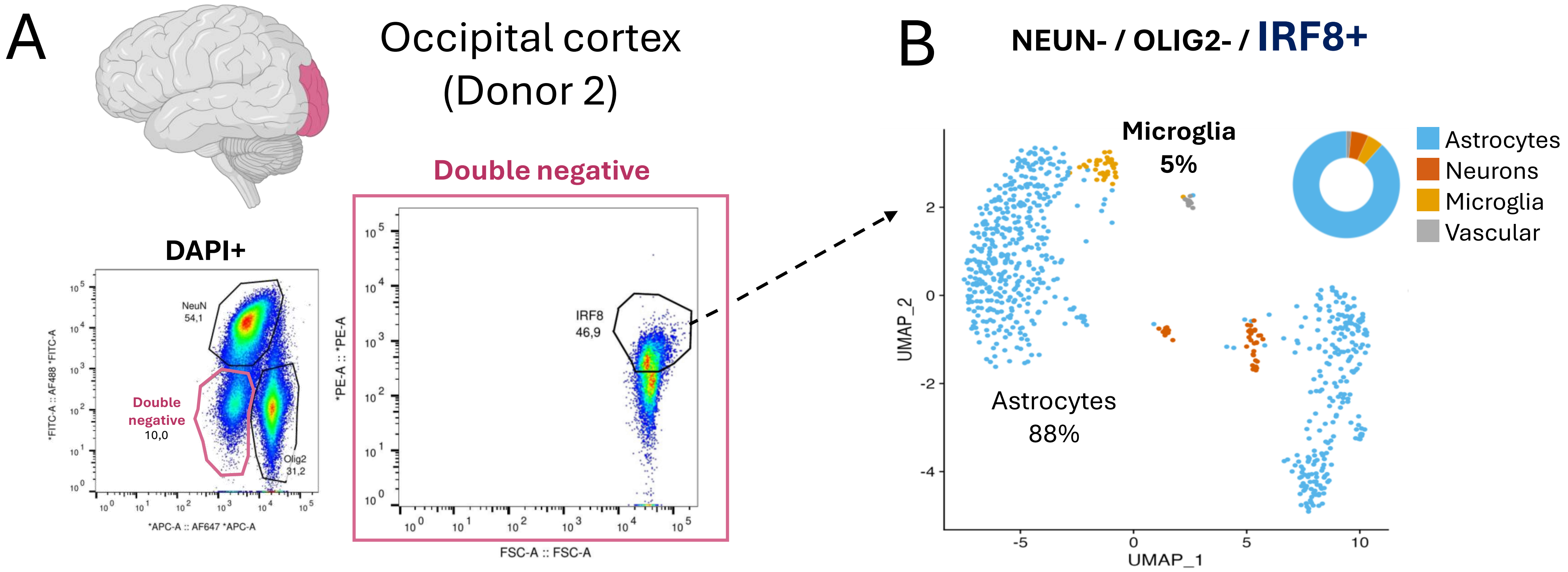


**Supplementary Figure 1. Cross-platform validation of the IRF8⁺ failure in occipital cortex.**

(A) Repeat of the IRF8$^+$ NEUN$^-$ OLIG2$^-$ sort on a separate Donor 2 occipital cortex aliquot (Methods Configuration A) processed by Illumina snPIP-seq library preparation rather than 10x Genomics (cf. Fig. 1C): hierarchical FACS gating tree (DAPI$^+$ → NEUN/OLIG2/Double-negative → IRF8$^+$ gate). (B) UMAP embedding and donut plot showing 88% astrocytes, 5% microglia and ~7% other (n = 760 nuclei). The reproducible recovery of an astrocyte-dominated population across two orthogonal sequencing platforms confirms that the IRF8$^+$ failure in occipital cortex reflects an intrinsic marker–region and/or donor limitation rather than a platform artefact.

Supplementary Figure 1

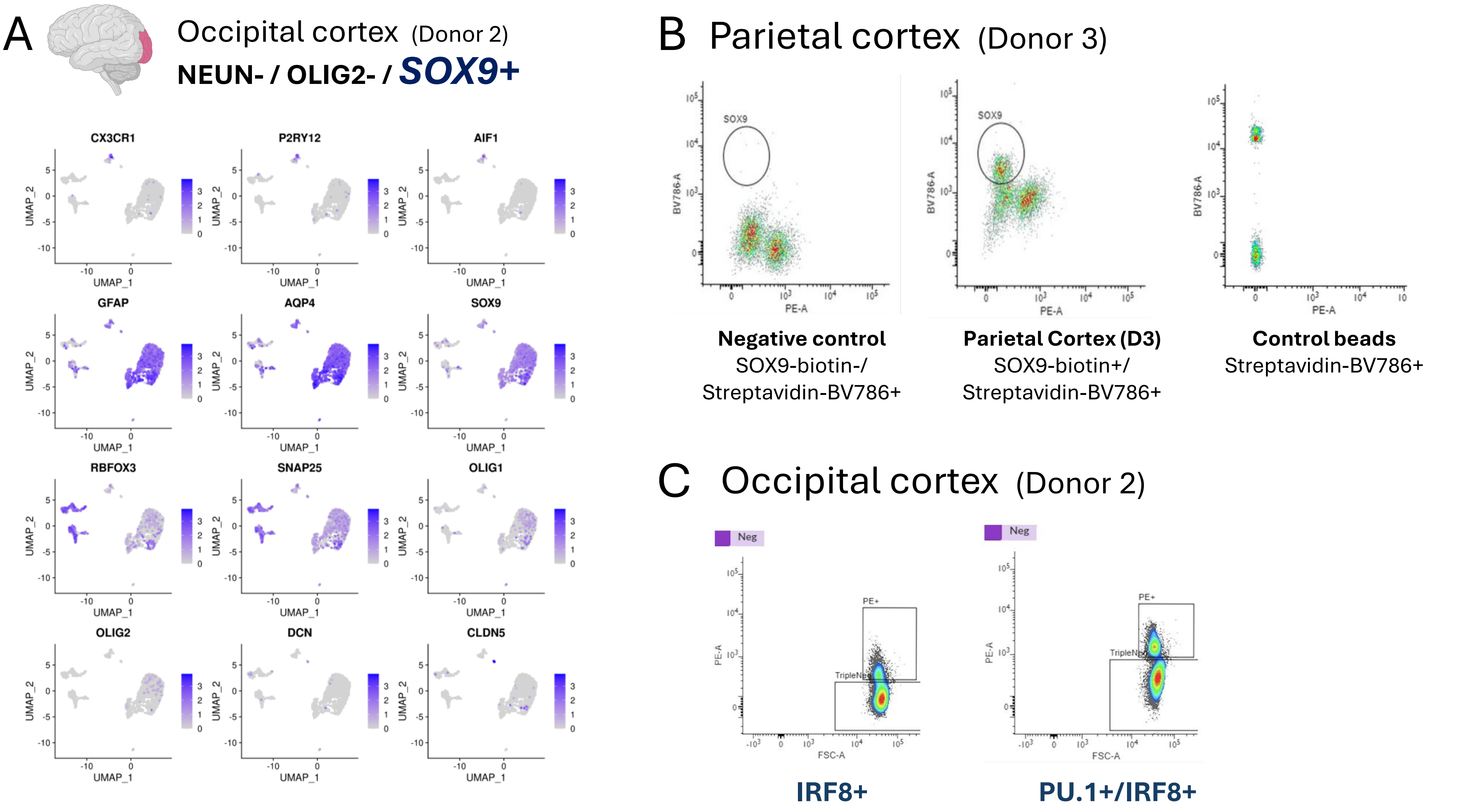


**Supplementary Figure 2. Validation of cell-type specificity, detection chemistry and dual-marker selection underlying the five-marker FANS panel.**

(A) Feature plots showing expression of canonical cell-type markers in the SOX9-PE enriched sample from Donor 2 occipital cortex: CX3CR1, P2RY12, AIF1 (microglia); GFAP, AQP4, SOX9 (astrocytes); RBFOX3, SNAP25 (neurons); OLIG1, OLIG2 (oligodendrocytes); DCN (fibroblast); CLDN5 (vascular). Expression values are log-normalised and all plots share the same scale. (B) Validation of the BV786-streptavidin detection system used for SOX9 in Methods Configuration D (Donor 3 parietal cortex): three side-by-side BV786-A versus PE-A scatter plots showing, from left to right, the negative control (sample stained without anti-SOX9 primary; SOX9-biotin$^{-}$ / Streptavidin-BV786$^{+}$), the stained sample (SOX9-biotin$^{+}$ / Streptavidin-BV786$^{+}$), in which a discrete SOX9$^{+}$ population is present, and BV786 control beads (instrument positive control). The selective appearance of the SOX9$^{+}$ population in the stained sample only confirms the specificity of the biotinylated anti-SOX9 / streptavidin-BV786 detection chain. (C) Comparison of IRF8-PE alone versus combined IRF8-PE + PU.1-PE staining on Donor 2 occipital cortex on the PE channel; the right plot shows improved separation of the PE$^{+}$ (microglia) and PE$^{-}$ (Triple-negative) populations with dual-marker selection on a shared PE channel.

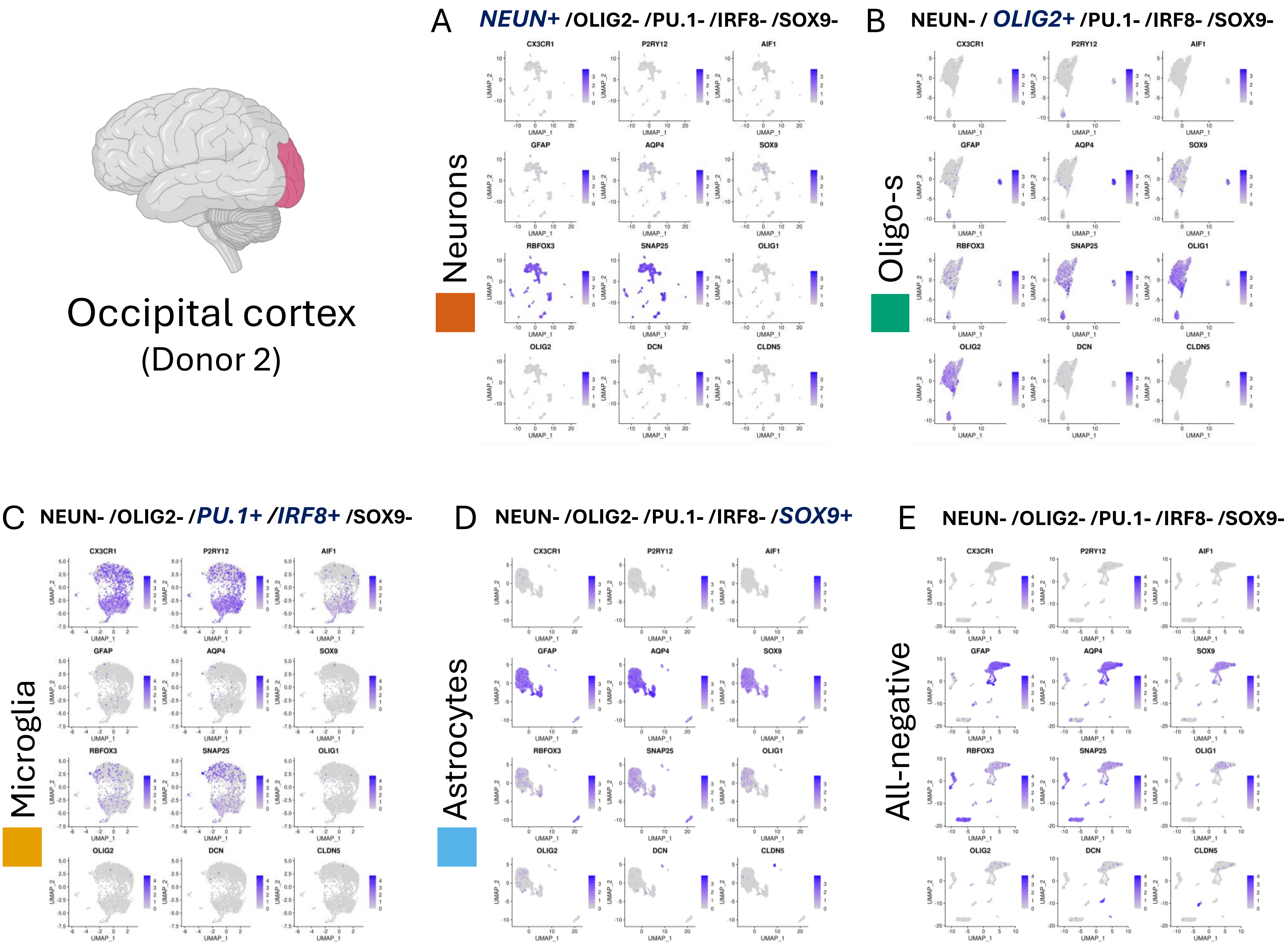


**Supplementary Figure 3. Canonical cell-type marker expression across the five sorted populations from the Configuration D five-marker sort (Donor 2 occipital cortex).**

Feature plots showing expression of canonical cell-type markers: CX3CR1, P2RY12, AIF1 (microglia); GFAP, AQP4, SOX9 (astrocytes); RBFOX3, SNAP25 (neurons); OLIG1, OLIG2 (oligodendrocytes); DCN (fibroblast); CLDN5 (vascular), on the UMAP embedding of each sorted population from Configuration D: (A) NEUN$^{+}$ neurons; (B) OLIG2$^{+}$ oligodendrocytes; (C) IRF8$^{+}$PU.1$^{+}$ microglia; (D) SOX9$^{+}$ astrocytes; (E) all-negative gate. Marker-expression patterns are consistent with the reference-anchored cell-type annotations reported in Figure 3. Expression values are log-normalised and all plots within each sorted population share the same scale.

Supplementary Figure 3

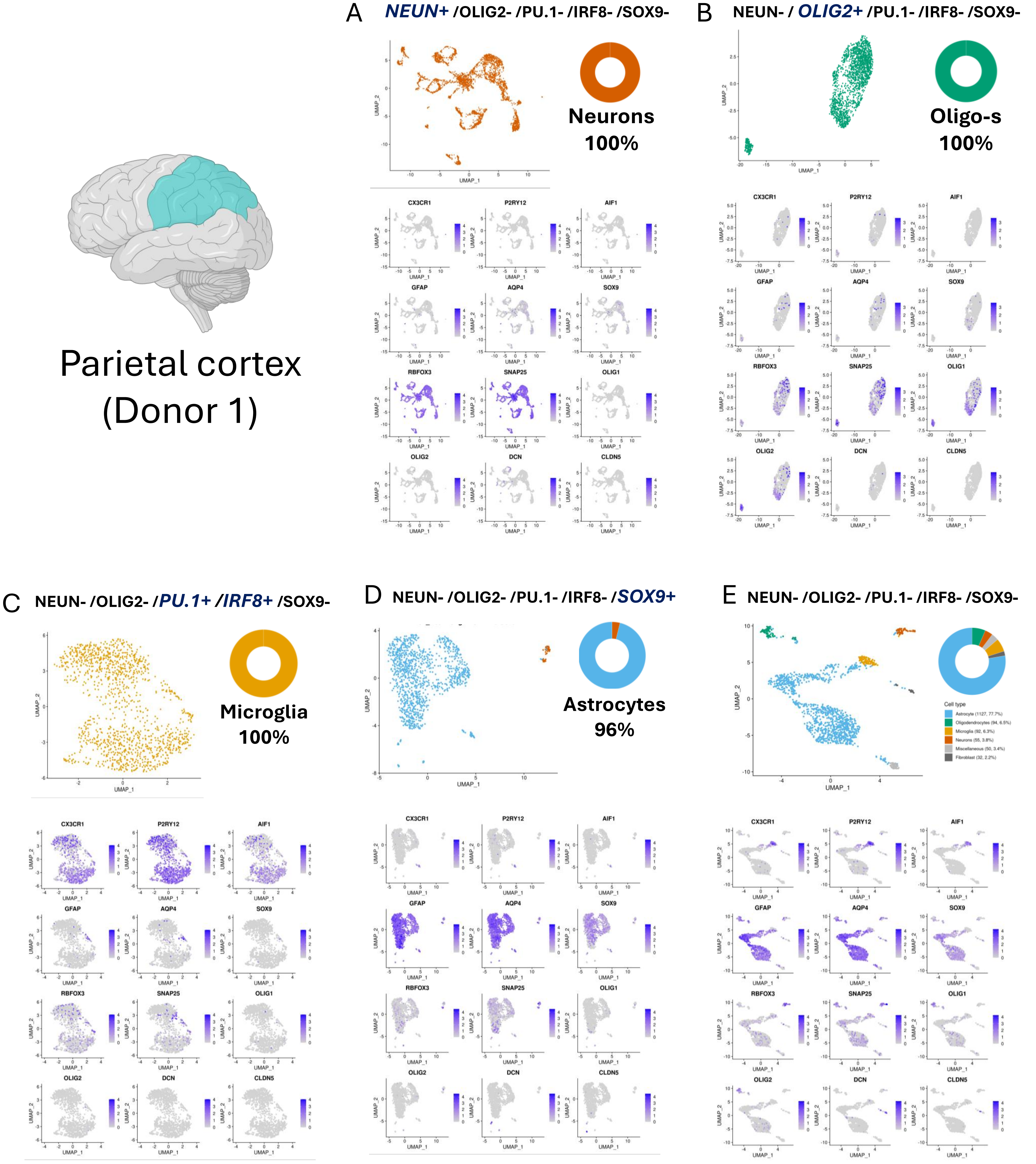

Parietal cortex
(Donor 1)
A NEUN+ /OLIG2- /PU.1- /IRF8- /SOX9-
Neurons
100%
B NEUN- / OLIG2+ /PU.1- /IRF8- /SOX9-
Oligo-s
100%
C NEUN- /OLIG2- /PU.1+ /IRF8+ /SOX9-
Microglia
100%
D NEUN- /OLIG2- /PU.1- /IRF8- /SOX9+
Astrocytes
96%
E NEUN- /OLIG2- /PU.1- /IRF8- /SOX9-
Cell type
Astrocyte (1127, 77.7%)
Oligodendrocytes (94, 6.5%)
Microglia (92, 6.3%)
Neurons (55, 3.8%)
Miscellaneous (50, 3.4%)
Fibroblast (32, 2.2%)
CX3CR1
P2RY12
AIF1
GFAP
AQP4
SOX9
RBFOX3
SNAP25
OLIG1
OLIG2
DCN
CLDN5
UMAP_1
UMAP_2

**Supplementary Figure 4. Cross-region replication of the five-marker panel in Donor 1 parietal cortex.**

Parallel five-population sort, repeated for another donor and cortical region (cf. Fig. 3, Supplementary Fig.3), reproduces the efficiency of the five-marker panel (Methods Configuration D; OYO-Link biotinylated anti-SOX9 detected with streptavidin-BV786). Feature plots showing expression of canonical cel
l-type markers: CX3CR1, P2RY12, AIF1 (microglia); GFAP, AQP4, SOX9 (astrocytes); RBFOX3, SNAP25 (neurons); OLIG1, OLIG2 (oligodendrocytes); DCN (fibroblast); CLDN5 (vascular), on the UMAP embedding of each sorted population from Configuration D applied to Donor 1 parietal cortex: (A) NEUN$^{+}$ neurons (100%; n=3,803); (B) OLIG2$^{+}$ oligodendrocytes/OPCs (100%; n=1,963); (C) IRF8$^{+}$PU.1$^{+}$ microglia (100%; n=1,286); (D) SOX9$^{+}$ astrocytes (96% astrocytes, 4% neurons; n= 1,208); (E) all-negative gate (77% astrocytes, 7% oligodendrocytes, 6% microglia, 4% neurons, 3% miscellaneous, 2% fibroblasts; n=1,450). Marker-expression patterns confirm the reference-anchored cell-type annotations depicted at the associated UMAPs. Expression values are log-normalised and all plots within each sorted population share the same scale.

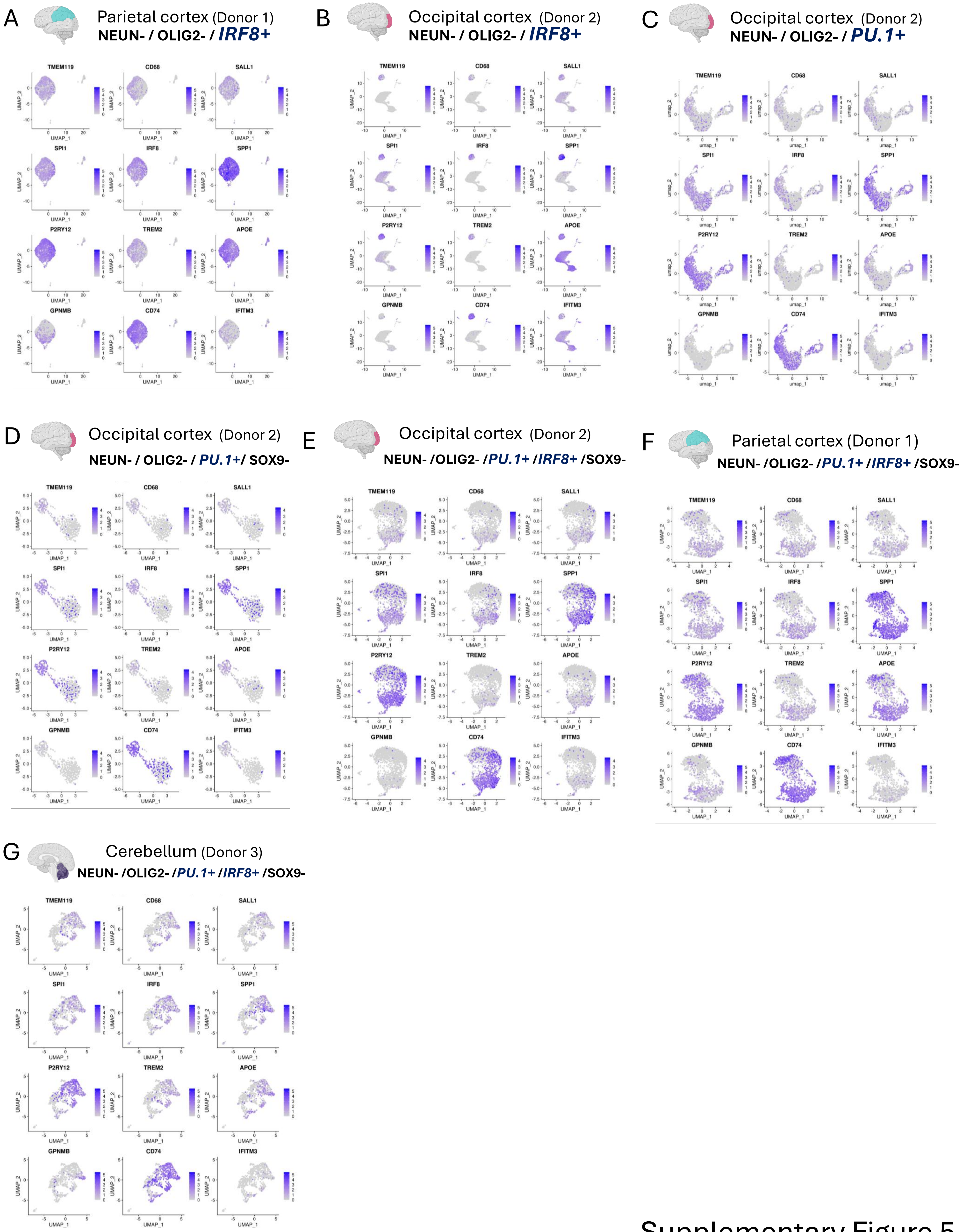


Supplementary Figure 5

**Supplementary Figure 5. Microglia subtype-marker expression across all microglia-enriched sorts.**

Feature plots showing expression of microglial homeostatic and activation-associated markers on the UMAP embedding of each microglia-targeted sort in the study: (A) Donor 1 parietal cortex IRF8$^+$ NEUN$^-$ OLIG2$^-$ (Methods Configuration A, 10x Genomics); (B) Donor 2 occipital cortex IRF8$^+$ NEUN$^-$ OLIG2$^-$ (Configuration A, 10x Genomics); (C) Donor 2 occipital cortex PU.1$^+$ NEUN$^-$ OLIG2$^-$ (Configuration A, snPIP-seq); (D) Donor 2 occipital cortex PU.1$^+$ SOX9$^-$ NEUN$^-$ OLIG2$^-$ (Configuration C, PU.1-only variant, snPIP-seq); (E) Donor 2 occipital cortex IRF8$^+$PU.1$^+$ five-marker sort (Configuration D, snPIP-seq); (F) Donor 1 parietal cortex IRF8$^+$PU.1$^+$ five-marker sort (Configuration D, snPIP-seq); (G) Donor 3 cerebellum IRF8$^+$PU.1$^+$ NEUN$^-$ OLIG2$^-$ SOX9$^-$ (Configuration C, IRF8+PU.1 variant, snPIP-seq). Expression values are log-normalised and all plots within each sorted population share the same scale.

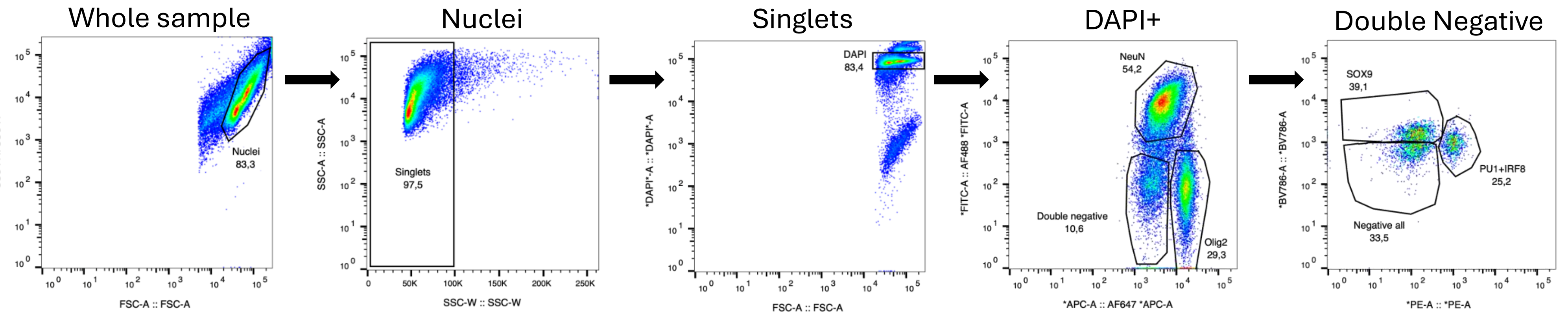


**Supplementary Figure 6. Hierarchical FANS gating strategy.**

Representative gating from a Donor 2 occipital cortex sort following Methods Configuration D. Plots are shown from left to right in the order of the gating hierarchy: (i) FSC-A × SSC-A "Nuclei" gate (83.3% of acquired events) excluding debris and aggregates; (ii) SSC-W × SSC-A "Singlets" gate (97.5%) excluding doublets; (iii) DAPI-A × FSC-A "DAPI$^+$" gate (83.4%) selecting intact nuclei from cellular debris; (iv) AF488 (NEUN) × AF647 (OLIG2) lineage plot resolving NEUN$^+$ neurons (54.2%), OLIG2$^+$ oligodendrocytes (29.3%) and the NEUN$^-$ OLIG2$^-$ "Double-negative" population (10.6%); (v) BV786 (SOX9) × PE (IRF8+PU.1) sub-gating of the Double-negative population into SOX9$^+$ astrocytes (39.1%), IRF8$^+$PU.1$^+$ microglia (25.2%) and an all-negative panel-defined fraction (33.5%). In Configurations A–C, the first four gate strategies are identical to those shown here, but the BV786 × PE sub-gating step (rightmost plot) is replaced by a PE-A × FSC-A plot used to draw the microglial-positive gate on the Double-negative population (see Methods).
